# Small Molecule Channels Harness Membrane Potential to Concentrate Potassium in trk1Δtrk2Δ Yeast

**DOI:** 10.1101/2020.04.06.028365

**Authors:** Jennifer Hou, Page N. Daniels, Martin D. Burke

## Abstract

Many protein ion channels harness membrane potential to move ions in opposition to their chemical gradient. Deficiencies of such proteins cause several human diseases, including cystic fibrosis, Bartter Syndrome Type II, and proximal renal tubular acidosis. Using yeast as a readily manipulated eukaryotic model system, we asked whether, in the context of a deficiency of such protein ion channel function *in vivo*, small molecule channels could similarly harness membrane potential to concentrate ions. In yeast, Trk potassium transporters use membrane potential to move potassium ions from a compartment of relatively low concentration outside cells (∼15mM) to one of >10 times higher concentration inside (150-500mM). trk1Δtrk2Δ yeast are missing these potassium transporters and thus cannot concentrate potassium or grow in standard media. Here we show that potassium permeable, but not potassium selective, small molecule ion channels formed by the natural product amphotericin B can harness membrane potential to concentrate potassium in trk1Δtrk2Δ cells and thereby restore growth. This finding expands the list of potential human channelopathies that might be addressed by a molecular prosthetics approach.

## INTRODUCTION

Many protein ion channels use membrane potential to move ions from compartments of relatively low concentration to those of higher concentration.^1^ These include the CFTR anion channel (which concentrates chloride), the ROMK potassium channel (which concentrates potassium), and the NBCe1 sodium bicarbonate cotransporter (which concentrates bicarbonate). Deficiencies of these protein functions cause the challenging to treat human diseases cystic fibrosis, Bartter Syndrome Type II, and proximal renal tubular acidosis, respectively.^2,3^ Replacing protein ion channels with small molecule mimics^4^ can in some cases restore physiology in cell and animal models of human diseases.^5,6,7^ With the goal of better understanding the potential scope and mechanistic underpinnings of this molecular prosthetics approach, we questioned whether small molecule channels, which tend to be less ion selective than their protein counterparts, could similarly harness membrane potential to concentrate certain ions in the context of a deficiency of protein ion channel function *in vivo*. We hypothesized that in such settings, increased membrane potential and/or compensatory actions of other protein transporters could enable imperfect small molecule ion channel mimics to be sufficient.

Yeast represent an excellent eukaryotic model system for asking such questions. In the plasma membrane of yeast, Trk1 and Trk2 potassium transporters^8,9,10^ harness membrane potential generated by the proton pump Pma1^11^ to move K^+^ from a compartment of relatively low concentration outside cells (∼15mM) to ≥10 times higher concentration inside cells (150-500mM), (Fig.1a).^12,13,14,15^ Deletion of Trk1 and Trk2 K^+^ transporters (trk1Δtrk2Δ)^12^ leads to decreased [K^+^]_intracellular_ (abbreviated [K^+^]_i_) and lack of cell growth under standard experimental conditions (Fig.1b).^12,16^ Interestingly, the plasma membranes of trk1Δtrk2Δ cells are hyperpolarized relative to wild type (Fig. S3),^12,13,17^ suggesting an amplified driving force for potassium import in the context of the potassium transporter deficiency. Moreover, there are a number of other protein transporters in yeast, including Tok1, Nha1, and Ena1^18^ that may be able to compensate for the relative lack of ion selectivity of a small molecule channel.

**Figure 1.**
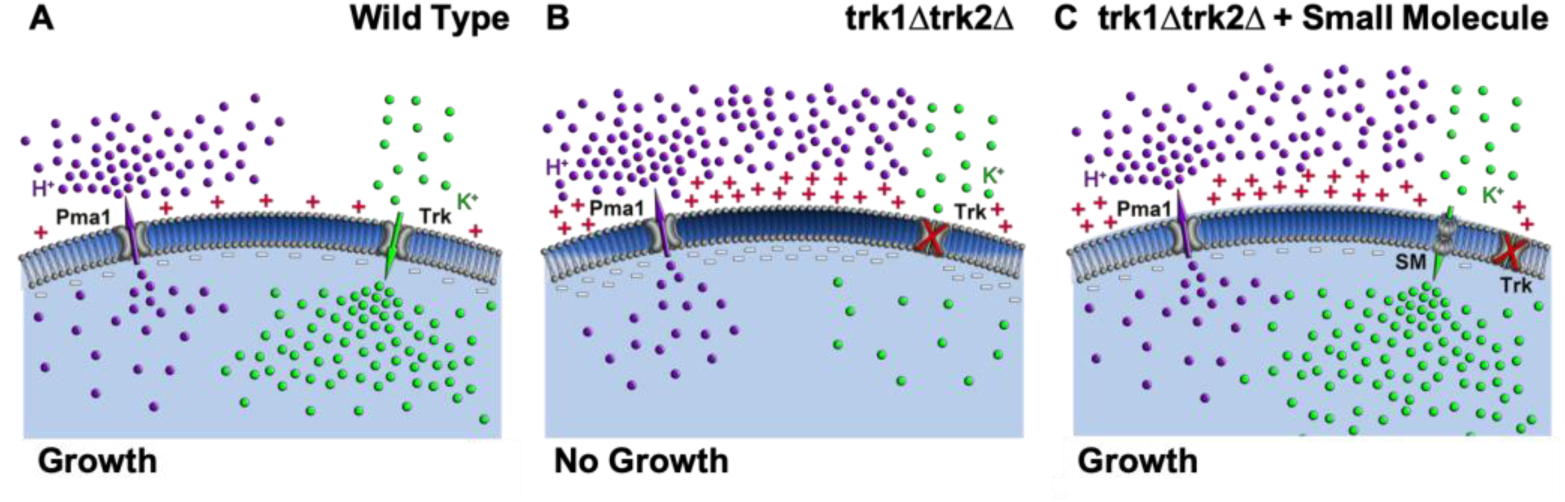
(a) Pma1 pumps H^+^ (purple spheres) out of cells and the resulting membrane potential is harnessed by Trk transporters to concentrate K^+^ (green spheres) inside cells. (b) trk1Δtrk2Δ yeast are hyperpolarized and yet unable to concentrate K^+^. (c) We hypothesized that small molecule mimics of the Trk potassium transporters can harness the membrane potential generated by Pma1 to concentrate K^+^ inside cells and thereby restore cell growth.

We recently reported that ion channels formed by the small molecule natural products amphotericin B (AmB) and nystatin A1 (Nyst) (Fig.S1) can restore growth in Trk potassium ion transporter-deficient yeast.^5^ We also noted that non-selective inhibition of Pma1 by ebselen blocks AmB- and Nyst-mediated growth rescue.^5^ However, in this prior study, we did not quantify intracellular potassium concentrations or relative membrane potential, nor did we have access to the recently reported more selective chemical inhibitor of Pma1 (Pma1_i_),^19^ thus limiting the mechanistic conclusions that could be made. To our knowledge, no prior studies determined whether small molecule channels can use membrane potential to concentrate ions. Here we test the mechanistic hypothesis that these imperfect small molecule mimics of Trk potassium ion transporters can harness the membrane potential in trk1Δtrk2Δ yeast to concentrate potassium inside these cells and thereby restore cell growth (Fig.1c).

## RESULTS

We first tested whether ion channels formed by AmB and Nyst^4^ could promote the net movement of K^+^ into trk1Δtrk2Δ yeast in opposition to its chemical gradient.^14,20^ In the absence of small molecule treatment, trk1Δtrk2Δ cells show reduced [K^+^]_i_ relative to WT cells (Fig.2a,c,e).^16^ Natamycin (Nat) is similar in structure to AmB and Nyst but does not form ion channels and thus served as a negative control.^21^ Nat did not change [K^+^]_i_ (Fig.2a), nor did it restore cell growth in trk1Δtrk2Δ at concentrations up to 1µM (Fig. 2b). In contrast, AmB caused a dose-dependent increase in [K^+^]_i_ in trk1Δtrk2Δ yeast (Fig.2c), which was paralleled by an increase in cell growth (Fig.2d). Notably, there was no change in [K^+^]_i_ or cell growth when WT cells were treated with the same low concentrations of AmB (Fig.2c,d). Similar results were observed with Nyst (Fig.2e,f).

**Figure 2.**
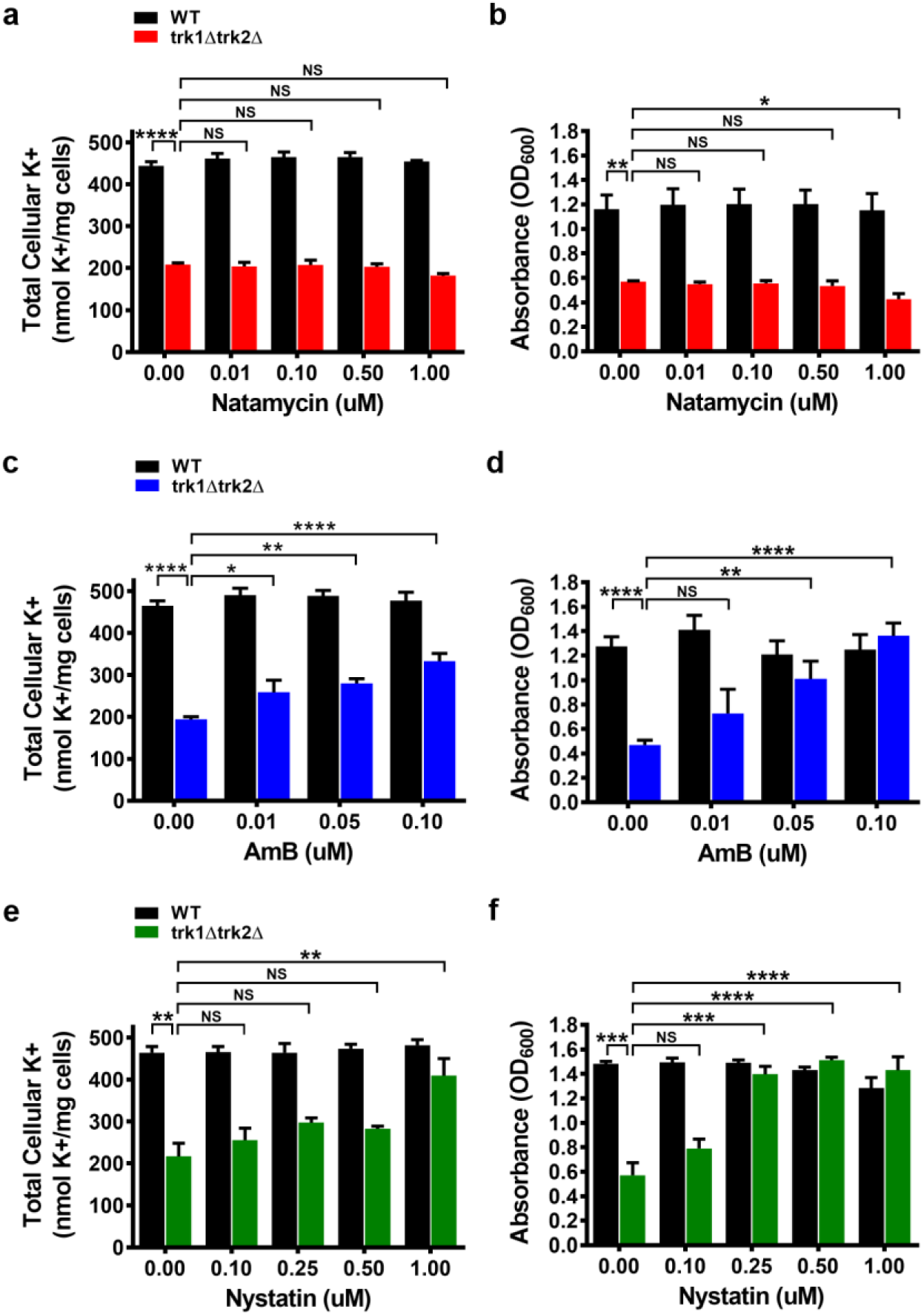
(a) Nat did not increase [K^+^]_i_, and (b) Nat did not rescue yeast growth. (c) AmB dose-dependently increased [K^+^]_i_ and (d) AmB similarly restored trk1Δtrk2Δ growth. (e) Nyst increased [K^+^]_i_ and (f) Nyst restored cell growth. *NS not significant*, \**P* ≤ 0.05, \*\**P* ≤ 0.01, \*\*\**P* ≤ 0.001, \*\*\*\**P* ≤ 0.0001.

We similarly quantified [K^+^]_i_ and cell growth as a function of time for a fixed concentration of each small molecule. We observed no changes in [K^+^]_i_ or cell growth over the course of 24h in trk1Δtrk2Δ cells treated with 100nM Nat (Fig.3a,b) compared to DMSO control. Similar results were observed when Nat was tested at 1µM (Fig.S2a,b). In contrast, 100nM AmB treatment increased [K^+^]_i_ to wild type-like levels within 6h (Fig.3c), and this was paralleled by WT-like growth rates (Fig.3d). Similar results were observed with 1µM Nyst (Fig.3e,f).

**Figure 3.**
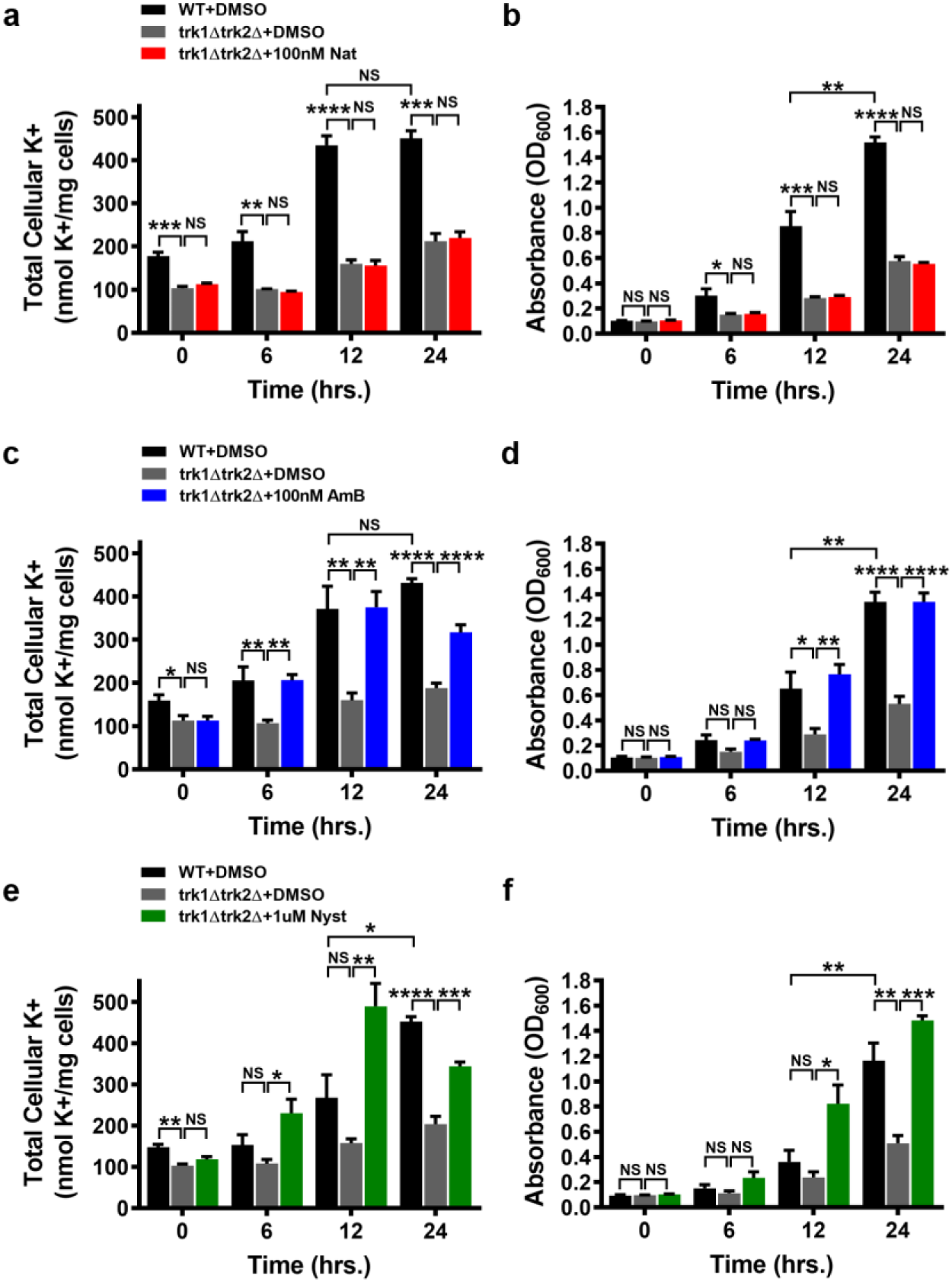
Small Molecule Ion Channels Increase [K^+^]_i_ over Time. Total cellular K^+^ content and growth were monitored in WT and small-molecule-treated trk1Δtrk2Δ yeast. (a) Nat did not increase [K^+^]_i_ (N=3) nor (b) rescue trk1Δtrk2Δ growth (N=3–4) when tested at 100nM, over the course of 24h. In contrast, both (c) AmB (100nM) and (e) Nyst (1µM) quickly restored [K^+^]_i_, and (d and f) rescued growth of trk1Δtrk2Δ cells (N=5 for (c),(d) and N=3 for (e),(f)). *NS not significant*, \**P* ≤ 0.05, \*\**P* ≤ 0.01, \*\*\**P* ≤ 0.001, \*\*\*\**P* ≤ 0.0001.

We next tested whether the AmB-mediated concentration of K^+^ ions in trk1Δtrk2Δ cells is driven by the membrane potential generated by Pma1. We employed the recently reported small molecule inhibitor of Pma1 (Pma1_i_)^19^ and flow cytometry to quantify the percentage of yeast cells that are relatively depolarized upon Pma1 inhibition. Cells with FITC-A signal above 10^3 were characterized as depolarized (to the right of black line, Fig.4a-c). In the absence of Pma1 inhibition a minimal percentage of trk1Δtrk2Δ cells were depolarized (<1%) (Fig. 4a,d). Modest inhibition of Pma1 by 15µM Pma1_i_ did not cause a statistically significant increase in the number of depolarized trk1Δtrk2Δ yeast cells (Fig. 4b,d). In contrast, most of the cells treated with 30µM Pma1_i_ were depolarized (78%, p≤ 0.001, Fig.4c,d).

**Figure 4.**
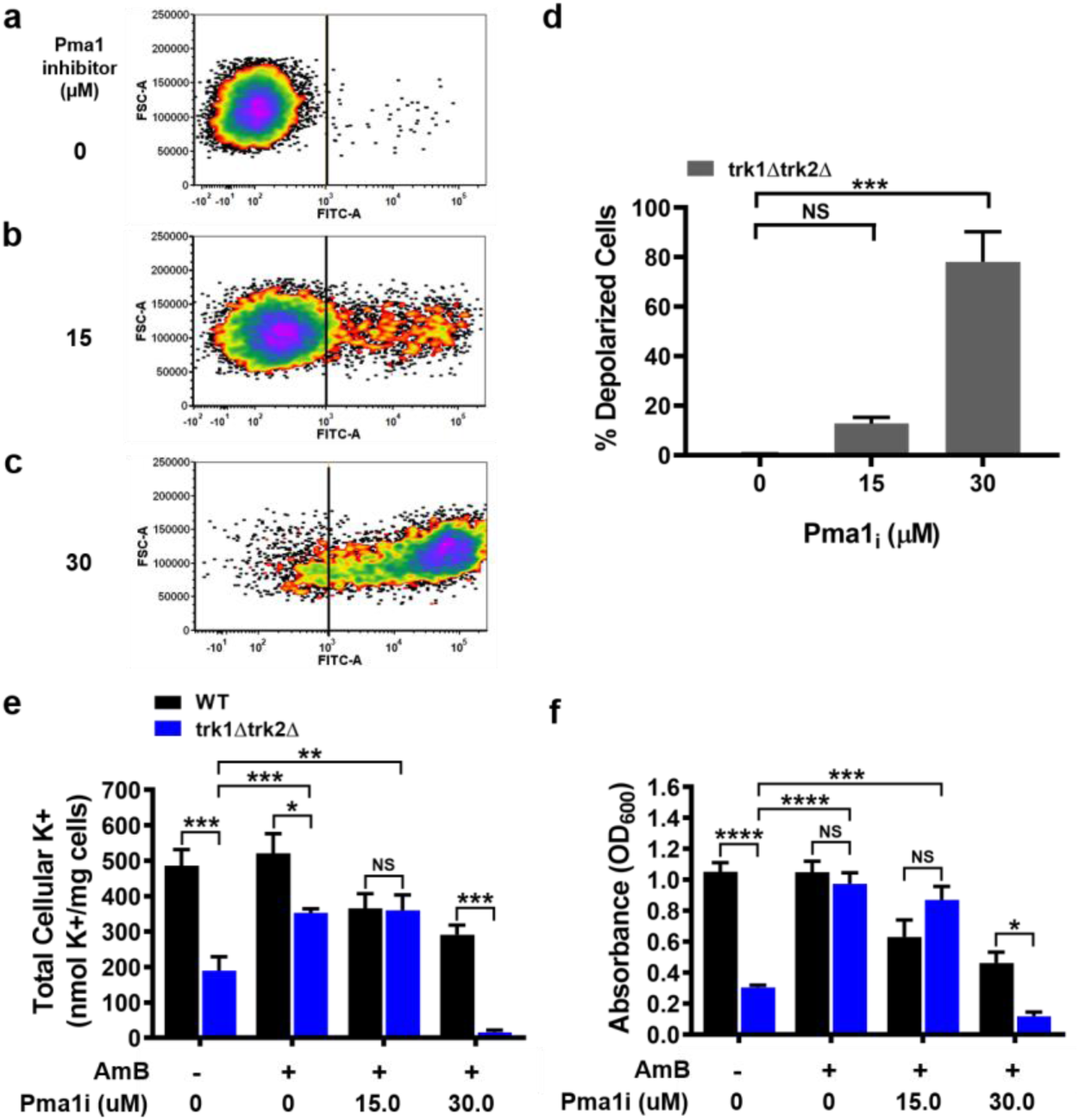
(a-c) Chemical inhibition with 30µM Pma1_i_ caused a substantial increase in the percentage of depolarized cells (N=3), representative flow cytometry data. (d) The corresponding % depolarized cells was determined as the percentage of cells for which cells with FITC-A signal above 10^3 were characterized as depolarized (right of black line), over the total number of live singlet cells (N=3). (e) Pma1 inhibition caused a reduction in total cellular K^+^ content. (f) Pma1 inhibition caused a reduction in cell growth in trk1Δtrk2Δ+100nM AmB relative to WT+100nM AmB cells. *NS not significant*, \**P* ≤ 0.05, \*\*\**P* ≤ 0.001, \*\*\*\**P* ≤ 0.0001.

Accordingly, trk1Δtrk2Δ yeast cells treated with 100nM AmB and either 0 or 15µM Pma1_i_ showed increases in [K^+^]_i_ (Fig.4e) and cell growth (Fig.4f) relative to non-AmB-treated trk1Δtrk2Δ yeast controls. In contrast, trk1Δtrk2Δ cells treated with 100nM AmB and 30µM Pma1_i_ showed no increase in [K^+^]_i_ nor cell growth (Fig.4e and f, respectively). AmB-treated trk1Δtrk2Δ yeast cells were also significantly more sensitive to this [Pma1_i_] than their similarly AmB-treated WT counterparts. Such disproportionate effects on [K^+^]_i_ and cell growth in trk1Δtrk2Δ and WT yeast were not observed when AmB-treated trk1Δtrk2Δ and WT yeast were alternatively treated with doxorubicin, which inhibits yeast growth via a non-ion channel-related mechanism (DNA intercalation) (Fig.S4). These results are consistent with the conclusion that Pma1-generated membrane potential drives the AmB-mediated concentration of K^+^ and restored cell growth in trk1Δtrk2Δ yeast.

## DISCUSSION

Thus, small molecule ion channels can harness membrane potential to concentrate ions inside cells and thereby restore physiology *in vivo*. These data suggest that the cellular mechanisms for concentrating K^+^ inside cells are sufficiently robust to enable such functional complementation with an imperfect small molecule mimic of the Trk proteins. It is also notable that AmB channels are not selective for K^+^; they can also transport Na^+^, H^+^, Cl^-^, and HCO_3_^-.6^ We speculate that other protein ion pumps and channels in the plasma membrane, such as Tok1, Nha1, and Ena1,^18^ may play a role in correcting for this imperfect ion selectivity, and this possibility will be interrogated in a future study. Collectively, these results further encourage the pursuit of small molecule surrogates for deficient protein ion channels that underlie a wide range of challenging to treat human diseases, including those that harness membrane potential to concentrate specific ions.

## METHODS

### Yeast Growth Conditions

Isogenic *Saccharomyces cerevisiae* strains BY4741(WT) and BYT12(trk1Δtrk2Δ) were maintained on 100mM KCl YPD agar plates at 4°C. Yeast were streaked every 1–3 weeks from frozen glycerol stocks. Overnight media was either 50 or 100mM KCl Yeast Nitrogen Base (YNB) pH 5.8 prepared by combining 4g (NH_4_)_2_SO_4_, 1.63g Translucent K^+^ Free YNB (ForMedium, CYN7505), 1.285g BSM (ForMedium, DBSM225), and the corresponding amount of KCl/L. pH of media was adjusted to 5.7 with citric acid and ammonium hydroxide before autoclaving. Following, 50mL of sterile 40% dextrose was added per liter. The same formulation was followed for K^+^ limiting media (15mM K^+^ final), with 1.11g of KCl added per liter.

### Determination of Relative Plasma Membrane Potential

This protocol was performed as described,^12^ with the following modifications. WT and trk1Δtrk2Δ cells were grown overnight in 50mM KCl supplemented YNB media, pH 5.8. After 18h, cells were harvested, washed, and re-suspended in 15mM KCl supplemented YNB, pH 5.8. UV-VIS spectrophotometry was used to determine the absorbance at 600nm (OD_600_). Cells were then diluted to an OD_600_=0.2 and incubated for indicated times. After incubation, cells were diluted to an OD_600_=0.2, if necessary. Each sample was pipetted into a 96-well black microplate. 3,3-dipropylthiacarbocyanine iodide (DiSC_3_(3)) dye was added to the wells, with a final concentration=0.2µM. Samples were incubated for 30min., 200XRPM, 30°C and then a fluorescence spectra, excitation 531nm, emission 549-589nm, was taken on a BioTek Synergy H1 microplate reader. The ratio of 580/560 was recorded for each sample.

### Flow Cytometry Analysis of Inhibition of Plasma Membrane H^+^-ATPase Pma1

Two to three colonies each of BY4741(WT) and BYT12(trk1Δtrk2Δ) were cultured overnight in 50mM KCl YNB pH 5.8, 200XRPM, 30°C. The following solutions were prepared before each experiment: 1mg/mL propidium iodide (PI) in H_2_O (final concentration=0.0033mg/mL) and 1.7mg/mL bis-(1,3 – dibutylbarbituric acid) trimethine oxonol (DiBAC) in DMSO (final concentration=0.001mg/mL)^19^ and sonicated. The following media were prepared: media (15mM KCl YNB pH 5.8, H_2_O, DMSO), PI media (15mM KCl YNB pH 5.8, PI stock, DMSO), DiBAC media (15mM KCl YNB pH 5.8, DiBAC stock, H_2_O), and PI and DiBAC media (15mM KCl YNB pH 5.8, PI stock, and DiBAC stock). 35mL of cells were pelleted at 1000XG, 23°C, for 5min. The supernatant was removed, and cells were resuspended in the corresponding volume of sterile milli-Q H_2_O. Then the cells were recentrifuged and washed. Afterwards, cells were resuspended in 15mM KCl YNB pH 5.8 to an OD_600_=0.090-0.110.

Next, 2mL of cell suspension were combined with the corresponding media condition. 6μL of DMSO or Pma1 inhibitor (Pma1_i_)^19^ stock solution was added to give the final concentrations of 15 and 30μM. Samples were incubated for 40-60min., 200XRPM, 30°C, and then analyzed on a BD LSRII Flow Cytometry Analyzer. Quality control samples were run before each experiment. Just before analyzing the sample, 30μL of AccuCheck counting beads were added to 270μL of sample and vortexed briefly. 2000 bead events were collected per sample. Afterwards, data was processed with FCS Express 7 software. First, cells were gated to exclude counting beads. Then based on PI staining intensity, dead cells were gated out (FCS-A vs. PI-A). The singlet cell population was analyzed, and cells with FITC-A signal above 10^3 were characterized as depolarized (to the right of black line, Fig. 4a-c). The same parameters were used to analyze data from three separate experiments.

### Small Molecule Rescue Assay of Potassium Transporter Deficient Yeast

All experiments were conducted under sterile conditions. Overnight yeast cultures comprised of three to five single colonies were grown to saturation over 12-24h, 200XRPM, 30°C in 100mM KCl YNB pH 5.8. Afterwards 90mL of cell suspension was pelleted at 1000XG, 23°C, for 5min. Supernatant was removed, and cells were resuspended in 90mL of sterile milli-Q H_2_O and vortexed. Cells were centrifuged and washed again. After removing the supernatant, cells were resuspended in 15mM KCl YNB, pH 5.8 and vortexed. Cells were diluted with 15mM KCl YNB media to OD_600_=0.090-0.110. Natamycin, Amphotericin B, and Nystatin A1 fresh stock solutions were prepared in DMSO and concentrations were determined by UV-VIS. UV-VIS specifications in methanol include Natamycin: λ_max_=317nm, ε=76,000M^-1^cm^-1^; Amphotericin B: λ_max_=406nm, ε=164,000M^-1^cm^-1^; Nystatin A1: λ_max_=304nm, ε=51,207 M^-1^cm^-1^. A 40-fold concentrated small molecule solution was prepared in DMSO for each treatment dose; and the small molecule solution was delivered as a 1:40 volume-to-volume ratio for small molecule to cell suspension volume. *Dose-response:* Per tested concentration, 100mL of cell suspension were treated with either DMSO control or small molecule. Cells were grown protected from light for 24h at 200XRPM, 30°C. *Time-course:* 500mL of cell suspension was treated with either DMSO or small molecule. Then 100mL of treated cell suspension was transferred into 250mL flasks designated for 6, 12, and 24h time points. The remaining 200mL per treatment group was designated for the 0h time point. Cells were cultured for each time point at 200XRPM, 30°C, protected from light. ^5, 12^

### Chemical Blocking of Small Molecule Mediated Rescue

The *Small Molecule Rescue Assay* protocol was modified to investigate the effect of chemical inhibition on protein ion pump Pma1. A 5µM AmB solution was prepared; the final ratio of AmB to cell suspension=1:50 and final ratio of chemical inhibitor to cell suspension=1:200. Doxorubicin and Pma1_i_ stock solutions were prepared in DMSO to achieve desired concentrations of 10mM and 6mM respectively. Subsequent doses were made by serial dilution.

Cells were harvested, washed, and diluted in 15mM KCl YNB, pH 5.8 media to OD_600_=0.090-0.110, as previously described. For DMSO controls, 2.5mL of DMSO were added to the respective 100mL each of WT+DMSO or trk1Δtrk2Δ+DMSO flasks. Separately, 500mL of WT and 500mL of trk1Δtrk2Δ cell suspensions were treated with 10mL each of 5µM AmB to give a final concentration of 100nM AmB. Then 100mL of each cell suspension was measured and transferred into labeled 250mL flasks. 0.5mL of DMSO or chemical inhibitor was added to each flask. The order of sample preparation was varied to avoid batch effects. Samples were cultured at 200XRPM, 30°C, 12h.

### Quantification of Total Cellular K^+^ Content via Inductively Coupled Plasma-Mass Spectrometry

At each time point, 1mL aliquots of each sample were utilized to determine OD_600_ values by UV-VIS. Subsequently, samples were quickly filtered through μm membrane filters (EMD Millipore) and washed with ice-cold 20mM MgCl_2_ (prepared with ultra-pure, molecule grade biology water). Filters were transferred into 6-well plates and 5mL of ice-cold 20mM MgCl_2_ was added to each well. Cells were resuspended; the resulting cell suspension was transferred into conical tubes and flash frozen in liquid nitrogen. Samples were lyophilized for ≥72h. Next, under argon, samples were transferred into aluminum capsules and sealed. Samples were analyzed by inductively coupled plasma-mass spectrometry.^16^

### Statistical Analysis

All statistical analyses were performed with GraphPad Prism software. Unpaired t-test with Gaussian distribution and one-way ANOVA with Dunnett’s multiple comparisons test were used for quantifying statistical differences between sample groups. *Statistical designations: NS not significant, *P ≤ 0*.*05, **P ≤ 0*.*01, ***P ≤ 0*.*001, and ****P ≤ 0*.*0001*.

## ASSOCIATED CONTENT

### Supporting Information

Supporting Information is available free of charge online.

Supporting data for transmembrane voltage and control studies.

## AUTHOR INFORMATION

### Notes

The University of Illinois has filed patent applications on small molecules that perform protein ion channel-like functions, and these have been licensed or optioned to Ambys Medicines and cystetic Medicines, companies for which M.D.B. is a founder and consultant.

## ACKNOWLEDGMENTS

We are grateful to Prof. Claudio Grosman for thoughtful review of this manuscript, Dr. Joaquín Ariño for the gift of the isogenic BY4741(WT) and BYT12(trk1Δtrk2Δ) *Saccharomyces cerevisiae* strains, Dr. Kiran Subedi, Elizabeth Eves, and Crislyn Lu for their assistance with inductively coupled plasma-mass spectrometry analysis of K^+^ content, which was performed at The UIUC Microanalysis Laboratory, and Dr. Barbara Pilas for her assistance with the flow cytometry protocol and analysis, which was performed at The Roy J. Carver Biotechnology Center at UIUC. We thank The National Institutes of Health (GM118185), and Howard Hughes Medical Institutes (HHMI) for funding support. P.N.D is an NIH Predoctoral Research Fellow (T32 GM070421). J.H. was supported partially by a UIUC Medical Scholars Program Fellowship.

## REFERENCES

(1) Gouaux, E.; MacKinnon, R. Principles of selective ion transport in channels and pumps. Science 2005, 310 (5753), 1461–1465.

(2) Jentsch, T.J.; Hübner, C.A.; Fuhrmann, J.C. Ion channels: function unraveled by dysfunction. Nat. Cell Biol. 2004, 6 (11), 1039–1047.

(3) (a) Ashcroft, F.M. From molecule to malady. Nature 2006, 440 (7083), 440–447. (b) Stoltz, D.A.; Meyerholz, D.K.; Welsh, M.J. Origins of Cystic Fibrosis Lung Disease. New Engl. J. Med. 2015, 372, 351–362. (c) Aronson, P.S.; Giebisch, G. Proximal renal tubular acidosis. In Genetic Disease of the Kidney; Lifton, R.P.; Somlo, S.; Giebisch, G.H.; Seldin, D.W., Eds.; New York, 2009; pp 199–212.

(4) (a) El-Etri, M.; Cuppoletti, J. Metalloporphyrin chloride ionophores: Induction of increased anion permeability in lung epithelial cells. Am. J. Physiol. 1996, 270, L386. (b) Busschaert, N.; Gale, P.A. Small-molecule lipid-bilayer anion transporters for biological applications. Angew. Chem., Int. Ed. 2013, 52, 1374. (c) Jiang, C.; Lee, E. R.; Lane, M. B.; Xiao, Y. F.; Harris, D. J.; Cheng, S. H. Partial correction of defective Cl-secretion in cystic fibrosis epithelial cells by an analog of squalamine. Am. J. Physiol. Lung Cell. Mol. Physiol. 2001, 281, L1164. (d) Koulov, A. V.; Lambert, T. N.; Shukla, R.; Jain, M.; Boon, J. M.; Smith, B. D.; Li, H.; Sheppard, D. N.; Joos, J. B.; Clare, J. P.; Davis, A. P. Chloride transport across vesicle and cell membranes by steroid-based receptors. Angew. Chem., Int. Ed. 2003, 42, 4931. (e) Shen, B.; Li, X.; Wang, F.; Yao, X.; Yang, D. A synthetic chloride channel restores chloride conductance in human cystic fibrosis epithelial cells. PLoS One 2012, 7, e34694. (f) Sakai, N.; Matile, S. Synthetic ion channels. Langmuir 2013, 29, 9031. (g) Fyles, T. M. How do amphiphiles form ion-conducting channels in membranes? Lessons from linear oligoesters. Acc. Chem. Res. 2013, 46, 2847. (h) Gokel, G. W.; Negin, S. Synthetic ion channels: From pores to biological applications. Acc. Chem. Res. 2013, 46, 2824.

(5) (a) Cioffi, A.G.; Hou, J.; Grillo, A.S.; Diaz, K.A.; Burke, M.D. Restored physiology in protein-deficient yeast by a small molecule channel. J. Am. Chem. Soc. 2015, 137 (32), 10096–10099. (b) In this previous study we found that AmB channels could increase the movement of ^86^Rb into yeast cells, but this experiment did not determine whether [K+]_i_ changed.

(6) (a) Muraglia, K.A.; Chorghade, R.S.; Kim, B.R.; Tang, X.X.; Shah, V.S.; Grillo, A.S.; Daniels, P.N.; Cioffi, A.G.; Karp, P.H.; Zhu, L.; Welsh, M.J.; Burke, M.D. Small-molecule ion channels increase host defense in cystic fibrosis airway epithelia. Nature 2019, 567 (7748), 405–408. (b) In this study we found that AmB channels could increase bicarbonate secretion and airway surface liquid pH in cultured CF airway epithelia, but in such settings bicarbonate is flowing down its concentration gradient. In contrast, CFTR uses membrane potential to concentrate chloride into the airway surface liquid, but chloride concentrations and apical membrane potential were not quantified in these studies.

(7) Grillo, A.S.; SantaMaria, A.M.; Kafina, M.D.; Cioffi, A.G.;, Huston, N.C.; Han, M.; Seo, Y.A.; Yien, Y.Y.; Nardone, C.; Menon, A.V.; Fan, J.; Svoboda, D.C.; Anderson, J.B.; Hong, J.D.; Nicolau, B.G.; Subedi, K.; Gewirth, A.A.; Wessling-Resnick, M.; Kim, J.; Paw, B.H.; Burke, M.D. Restored iron transport by a small molecule promotes absorption and hemoglobinization in animals. Science 2017, 356 (6338), 608–616.

(8) Gaber, R.F.; Styles, C.A.; Fink, G.R. TRK1 encodes a plasma membrane protein required for high-affinity potassium transport in Saccharomyces cerevisiae. Mol. Cell. Biol. 1988, 8 (7), 2848–2859.

(9) Ko, C.H.; Buckley, A.M.; Gaber, R.F. TRK2 is required for low affinity K+ transport in Saccharomyces cerevisiae. Genetics 1990, 125 (2), 305–312.

(10) Ko, C.H.; Gaber, R.F. TRK1 and TRK2 encode structurally related K+ transporters in Saccharomyces cerevisiae. Mol. Cell. Biol. 1991, 11 (8), 4266–4273.

(11) (a) Andre, B. An overview of membrane transport proteins in Saccharomyces cerevisiae. Yeast 1995, 11 (16), 1575–1611. (b) Ariño, J.; Ramos, J.; Sychrová, H. Alkali metal cation transport and homeostasis in yeast. Microbiol. Mol. Biol. Rev. 2010, 74 (1), 95–120.

(12) Navarrete, C.; Petrezsélyová, S.; Barreto, L.; Martinez, J.L.; Zahrádka, J.; Ariño, J.; Sychrová, H.; Ramos, J. Lack of main K+ uptake systems in Saccharomyces cerevisiae affects yeast performance in both potassium-sufficient and potassium-limiting conditions. FEMS Yeast Res. 2010, 10 (5), 508–517.

(13) Zahrádka, J.; Sychrová, H. Plasma-membrane hyperpolarization diminishes the cation efflux via Nha1 antiporter and Ena ATPase under potassium-limiting conditions. FEMS Yeast Res. 2012, 12 (4), 439–446.

(14) Cyert, M.S.; Philpott, C.C. Regulation of cation balance in Saccharomyces cerevisiae. Genetics 2013, 193 (3), 677–713.

(15) Herrera, R.; Álvarez, M.C.; Gelis, S.; Ramos, J. Subcellular potassium and sodium distribution in Saccharomyces cerevisiae wild-type and vacuolar mutants. Biochem. J. 2013, 454 (3), 525–532.

(16) Gelis, S.; González-Fernández, R.; Herrera, R.; Jorrín, J.; Ramos, J. A physiological, biochemical and proteomic characterization of Saccharomyces cerevisiae trk1,2 transport mutants grown under limiting potassium conditions. Microbiology 2015, 161 (6), 1260–1270.

(17) Herrera, R.; Álvarez, M.C.; Gelis, S.; Kodedová, M.; Sychrová, H.; Kschischo, M.; Ramos, J. Role of Saccharomyces cerevisiae Trk1 in stabilization of intracellular potassium content upon changes in external potassium levels. Biochim. Biophys. Acta. 2014, 1838 (1), 127–133.

(18) Volkov, V. Quantitative description of ion transport via plasma membrane of yeast and small cells. Front. Plant Sci. 2015, 6 (425), 1–20.

(19) Kjellerup, L.; Gordon, S.; Cohrt, K.O.; Brown, W.D.; Fuglsang, A.T.; Winther, A.M.L. Identification of antifungal H+-ATPase inhibitors with effect on plasma membrane potential. Antimicrob. Agents Chemother. 2017, 61 (7), e00032–17.

(20) Ariño, J.; Ramos, J.; Sychrová, H. Monovalent cation transporters at the plasma membrane in yeasts. Yeast 2018, 36 (4), 177–193.

(21) Welscher, Y.M.; ten Napel, H.H.; Balagué, M.M.; Souza, C.M.; Riezman, H.; de Kruijff, B.; Breukink, E. Natamycin blocks fungal growth by binding specifically to ergosterol without permeabilizing the membrane. J. Biol. Chem. 2008, 283 (10), 6393–6401.

