## Supplemental Info for "Small Molecule Channels Harness Membrane Potential to Concentrate Potassium in trk1Δtrk2Δ Yeast"

**Table of Contents**

**S1** Fig. S1. Chemical Structures of Small Molecule Probes.
**S2** Fig. S2. Channel-Inactive-Small-Molecule Natamycin Does Not Restore $K^+$ Content nor Cell Growth in Protein-Deficient Yeast.
**S3** Fig. S3. A Transmembrane Voltage Exists in Potassium Transporter Deficient Yeast under Conditions of $K^+$ Limitation.
**S4** Fig. S4. Doxorubicin Similarly Reduces Growth in AmB Treated WT and trk1Δtrk2Δ Yeast.

### Supplemental Figures

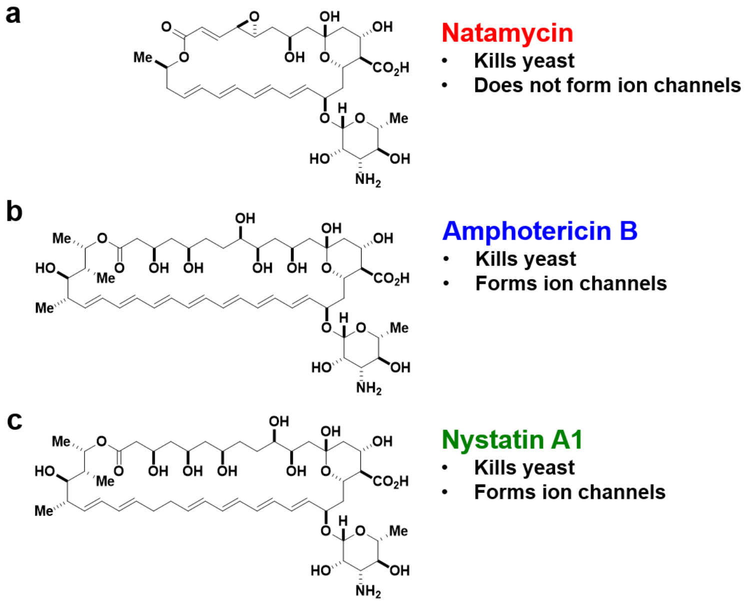


**Fig. S1.** Chemical Structures of Small Molecule Probes. (a) Natamycin (Nat), (b) Amphotericin B (AmB), and (c) Nystatin A1 (Nyst).

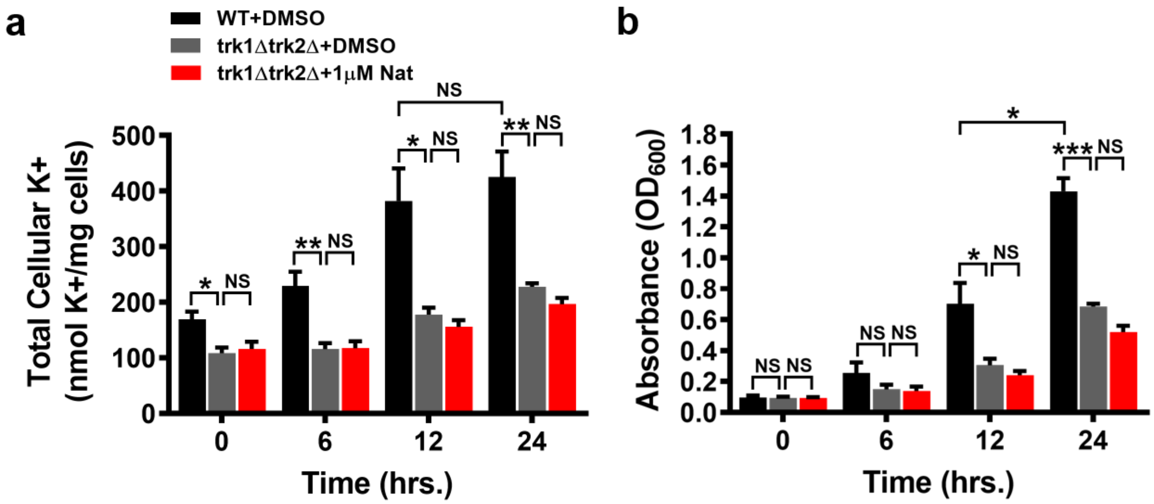


**Fig. S2.** Channel-Inactive-Small-Molecule Natamycin Does Not Restore $K^+$ Content nor Cell Growth in Protein-Deficient Yeast. 1μM Nat did not increase total cellular $K^+$ content (a) nor promote trk1Δtrk2Δ cell growth (b) across all time points tested (6, 12, and 24h), N=3, *NS not significant,* *$P \le 0.05$, **$P \le 0.01$, ***$P \le 0.001$.

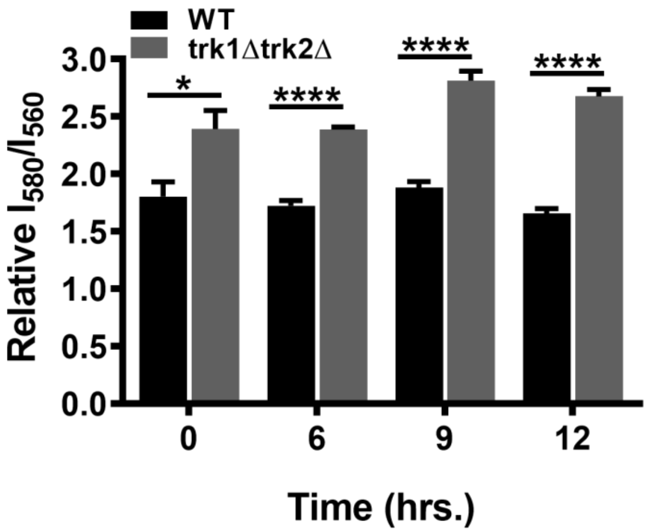


**Fig. S3.** A Transmembrane Voltage Exists in Potassium Transporter Deficient Yeast under Conditions of $K^+$ Limitation. To assess the relative plasma membrane potential, WT and trk1Δtrk2Δ yeast were transferred from 50mM to 15mM KCl containing media and incubated for the indicated times before adding DiSC3(3) dye. Fluorescence ratio ($I_{580}/I_{560}$) showed that trk1Δtrk2Δ cells were hyperpolarized compared to WT at 0, 6, 9, and 12h. N=6 samples for 0h, N≥11 samples for 6, 9, and 12h, over three independent experiments. **P* ≤ 0.05, ***P* ≤ 0.01, *****P* ≤ 0.0001.

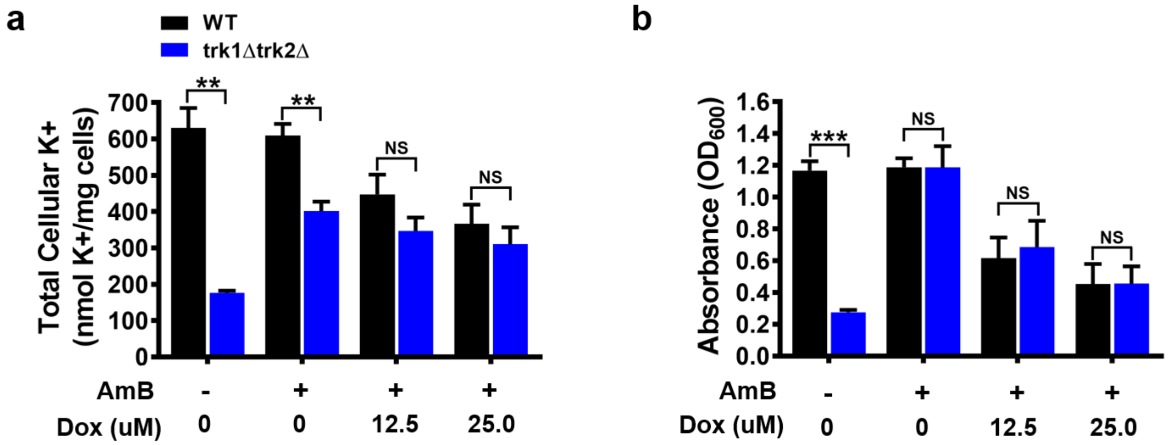


**Fig. S4.** Doxorubicin Similarly Reduces Growth in AmB Treated WT and trk1Δtrk2Δ Yeast. (a) At 12h, Doxorubicin, a chemical inhibitor that blocks cell growth by a different mechanism (DNA intercalation which is unrelated to ion transport) caused minor losses in $K^+$ content in WT+100nM AmB and trk1Δtrk2Δ+100nM AmB. (b) Furthermore, both treatment groups showed similar diminishment of cell growth. N=3, *NS not significant*, ***P* ≤ 0.01, ****P* ≤ 0.001.
